# Multiple binding modes of ibuprofen in human serum albumin identified by absolute binding free energy calculations

**DOI:** 10.1101/068502

**Authors:** Stefania Evoli, David L. Mobley, Rita Guzzi, Bruno Rizzuti

## Abstract

Human serum albumin possesses multiple binding sites and transports a wide range of ligands that include the anti-inflammatory drug ibuprofen. A complete map of the binding sites of ibuprofen in albumin is difficult to obtain in traditional experiments, because of the structural adaptability of this protein in accommodating small ligands. In this work, we provide a set of predictions covering the geometry, affinity of binding and protonation state for the pharmaceutically most active form (S– isomer) of ibuprofen to albumin, by using absolute binding free energy calculations in combination with classical molecular dynamics (MD) simulations and molecular docking. The most favorable binding modes correctly reproduce several experimentally identified binding locations, which include the two Sudlow’s drug sites (DS2 and DS1) and the fatty acid binding sites 6 and 2 (FA6 and FA2). Previously unknown details of the binding conformations were revealed for some of them, and formerly undetected binding modes were found in other protein sites. The calculated binding affinities exhibit trends which seem to agree with the available experimental data, and drastically degrade when the ligand is modeled in a protonated (neutral) state, indicating that ibuprofen associates with albumin preferentially in its charged form. These findings provide a detailed description of the binding of ibuprofen, help to explain a wide range of results reported in the literature in the last decades, and demonstrate the possibility of using simulation methods to predict ligand binding to albumin.

**Graphical abstract:** 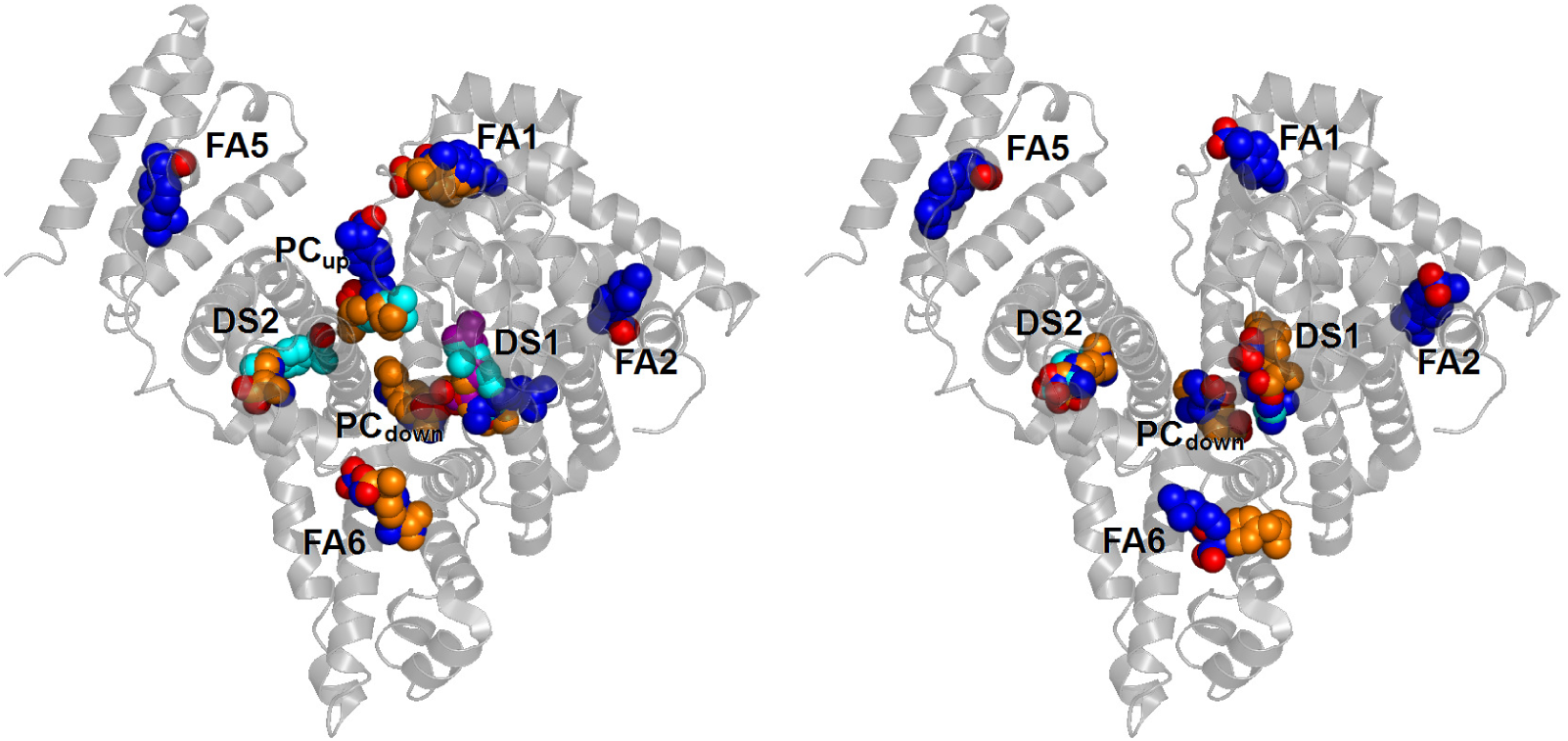

**Focus:** Alchemical free energy methods can identify favored binding modes of a ligand within a large protein with multiple binding sites

**Highlights:**

- Human serum albumin binds the anti-inflammatory drug ibuprofen in multiple sites
- Alchemical free energy calculations predicted favored binding modes of ibuprofen
- Bound geometry, affinity and protonation state of the ligand were determined
- Simulations identified a number of previously undetected binding sites for ibuprofen
- Free energy methods can be used to study large proteins with multiple binding sites

## 1. Introduction

Human serum albumin is the most abundant protein in blood plasma, where it acts as a carrier of non-esterified fatty acids and several other endogenous and exogenous compounds, including a large variety of pharmaceuticals and active metabolites (Fasano et al. 2005).

Albumin is monomeric (585 amino acid residues, 66 kDa), with a predominantly α-helical secondary content and a heart-shaped tertiary conformation (Curry et al. 1998). The protein structure encompasses three homologous domains (I-III), each divided in two sub-domains (A-B) connected through random coils. Long-chain fatty acids, as well as many other different ligands, attach to albumin in seven binding pockets with corresponding names FA1 through FA7 (Bhattacharya et al. 2000). Site FA7 can enlarge to accommodate ligands bulkier than fatty acids, extending towards site FA2 to form Sudlow’s drug site 1 (Sudlow et al. 1975), or DS1. Similarly, the two sites FA3 and FA4 can rearrange into a single and larger drug site, DS2. Other binding sites for complex heterocyclic drugs have also been detected or proposed (Handing et al. 2016; Wang et al. 2013). Some smaller ligands can additionally bind along the protein cleft in between domains IB-IIIA, or in an extension of site FA2 completely included in domain IA, called site FA2ʹ when occupied by lauric or myristic acid (Bhattacharya et al. 2000). Overall, the binding features of albumin are due to such a multiplicity of available binding pockets, combined with its intrinsic structural plasticity in accommodating several types of compounds.

Ibuprofen is a non-steroidal drug (Fig. 1) used for its anti-inflammatory effects, which has long been considered as stereotypical ligand for site DS2 (Sudlow et al. 1975). It is almost insoluble in water, and in ordinary therapeutic quantity is 99% bound and carried in blood plasma through proteins (Fanali et al. 2012), mostly by human serum albumin. The carboxyl group of ibuprofen has p*K*_a_ ≈ 4.4 (Babic et al. 2007), thus it is expected to be deprotonated (−COO^−^) under physiological conditions (pH of blood is 7.40 ± 0.05). However, within the core of a protein matrix the pH may easily differ by up to three units compared to the surrounding solvent (Onufriev and Alexov 2013). Thus, ibuprofen might be complexed with albumin in either or both neutral or charged states, and the assumption that it binds only in deprotonated form needs to be verified. In addition, due to the presence of a chiral center involving the central carbon atom in the propionic acid moiety (see Fig. 1), ibuprofen has two enantiomers. In commercial formulations, the drug is commonly available in a racemic mixture of both isomers. However, only S–ibuprofen is pharmacologically active, as demonstrated in both *in vitro* and *in vivo* studies (Cheng et al. 1994; Paliwal et al. 1993). Furthermore, the R– isomer largely interconverts into S–ibuprofen as a result of metabolism (Cheng et al. 1994).

**Fig. 1.**
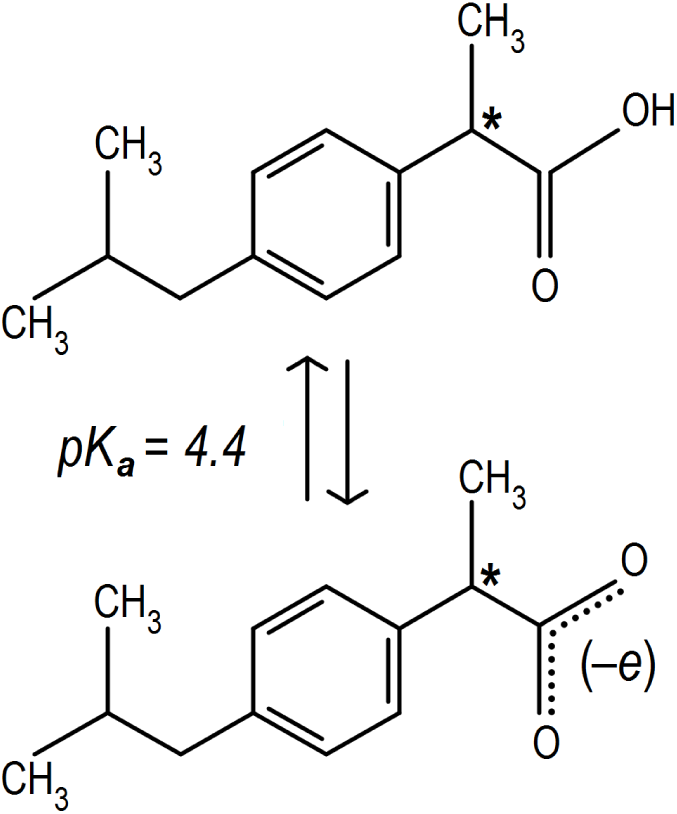
Structure of ibuprofen: (top) protonated neutral, and (bottom) deprotonated charged form. Molecular mass is, respectively, 206.3 and 205.3 Da for the two forms. The only chiral C atom is indicated with an asterisk.

Ibuprofen shows a very high affinity for albumin, with measured binding constants up to 2⋅10^6^ M^−1^ (Itoh et al. 1997; Rahman et al. 1993), and primarily binds in site DS2, as demonstrated by competition for binding with the marker ligand diazepam (Kosa et al. 1997). The crystallographic structure of the molecular complex of albumin with S–ibuprofen (Ghuman et al. 2005) confirmed this binding location and demonstrated the presence of a secondary site corresponding to FA6. The ligand is assumed to be deprotonated in both cases, although no direct evidence is available because hydrogen atoms are not resolved in the crystal. Ibuprofen was also found to bind with lower affinity in the DS1 site, but the observed electron density was too weak to define the conformation and orientation of a binding pose (Ghuman et al. 2005). Another weak binding location has been proposed in site FA2 (di Masi et al. 2011), but the exact anchoring geometry and protonation of ibuprofen is unknown also in this case. Several other spectroscopic techniques suggested the presence of possible, additional secondary sites in unidentified locations: the estimate of the binding parameters of ibuprofen to albumin by equilibrium dialysis (Kosa et al. 1997) required a total of 6 sites to fit the experimental data, whereas up to 10 binding modes were suggested in other cases (Montero et al. 1990; Whitlam et al. 1979).

Molecular dynamics (MD) simulations have already been used in several works to elucidate details of the binding location, conformation and affinity of ligands associated with human serum albumin (Deeb et al. 2010; Fujiwara and Amisaki 2008; Pantusa et al. 2014; Rizzuti et al. 2015). Alchemical free energy methods are currently emerging as one of the most powerful tools to study the binding of drugs to host proteins (Chodera et al. 2011; Christ and Fox 2014; Gallicchio and Levy 2011; Gumbart et al. 2013; Michel 2014; Mobley and Klimovich 2012; Wang et al. 2015) but (to the best of our knowledge) have not been applied so far to a large protein with multiple binding sites, such as albumin. This is partly because, among alchemical calculations, relative free energy calculations are not suitable for predicting binding sites or comparing binding of the same ligand in multiple sites, and absolute free energy calculations have not yet seen much mainstream usage. In this work, by using a combination of molecular docking, conventional MD simulations and absolute alchemical free energy calculations, we report predicted bound geometries, protonation state and affinity for the binding of the pharmaceutically active form of ibuprofen (S–ibuprofen) to human serum albumin. The results are in excellent agreement with previous experimental observations, appear to exclude the possibility that ibuprofen binds the protein in neutral form, and uncover several previously undetected binding modes, ultimately demonstrating that simulation techniques can reveal significant details of the binding properties of albumin.

## 2. Computational methods

### 2.1 Molecular modeling and docking

The structure of unliganded albumin was obtained from the complex crystallized in the presence of two ibuprofen molecules (Ghuman et al. 2005), deposited as 2BXG entry in the Protein Data Bank. The position of seven missing residues at the two protein backbone termini, as well as missing atoms in solvent-exposed side chains of some other residues, was reconstructed *in silico* by using VMD (Humphrey et al. 1996).

Molecular docking of S–ibuprofen to albumin was carried out through AutoDock Vina (Trott and Olson 2010). The graphical interface AutoDock Tools 1.5.6 (Morris et al. 1998) was preliminarily used to add polar hydrogen atoms to the protein, determine the allowed torsions for the ligand, and define the coordinates for the search space.

The docking procedure consisted of 12 independent runs, each determining the best 20 docking conformations ranked according to their binding affinity. The search space included the entire protein volume in an unbiased way. The resulting 240 poses were reduced to 38 through a clustering procedure based on distances, that included binding modes with a RMSD (Root Mean Square Deviation) of atomic positions < 2.5 Å in each cluster.

### 2.2 Molecular dynamics simulations

MD simulations were performed for albumin complexed with a single ibuprofen molecule, either neutral or charged, placed in each of the 38 binding modes previously selected, for a total of 76 runs. The simulation package GROMACS 4.6.3 (Pronk et al. 2013) was used in combination with the AMBER 99SB-ILDN force field for the protein (Lindorff-Larsen et al. 2010) and GAFF for the ligand (Wang et al. 2004). The topologies for ibuprofen was built by using AmberTools 13 (Wang et al. 2006) and atomic charges were assigned by using the AM1-BCC method (Jakalian et al. 2002), as implemented in the Antechamber module.

Protein residues were adapted to mimic a physiological pH, with positively charged Arg/Lys side chains and N-terminus, and negatively charged Asp/Glu side chains and C-terminus. The complex was placed in a rhombic dodecahedron box with a minimum distance of 1 nm to the nearest box edge, and surrounded with ~ 32000 water molecules described by the TIP3P water model (Jorgensen et al. 1983). The overall charge of the system was neutralized by adding 15 Na^+^ counterions in simulations with neutral ibuprofen, and 16 for the charged form. Periodic boundary conditions were applied to avoid edge effects. The energy of the system was minimized by using a steepest descendent method for 1500 steps.

Initial velocities were obtained from a Maxwell-Boltzmann distribution at 300 K, and the system was equilibrated for 10 ps at constant volume and temperature. Production runs were carried out for 5 ns in the NPT ensemble, using a Berendsen barostat with a time constant 0.5 ps to control the pressure (Berendsen et al. 1984) and a Langevin thermostat with inverse friction coefficient 0.1 ps. The electrostatic interactions were treated with the Particle-Mesh Ewald (PME) method (Darden et al. 1993; Essmann et al. 1995), with 1.2 nm cut-off between direct and reciprocal space summation, and 6^th^ order spline interpolation with a grid spacing of 0.1 nm. The van der Waals interactions were modeled with a switched Lennard-Jones potential cut-off at 1 nm, with long range dispersion corrections applied for both energy and pressure. Bond distances involving hydrogen atoms were constrained with the P-LINCS algorithm (Hess 2008) and an integration time step of 2 fs was used.

### 2.3 Clustering and selection of binding modes

Binding modes of ibuprofen that were stable during the MD simulations were selected on the basis of their position and predicted binding and/or interaction energy. To this aim, the configurations of the complex in the individual MD trajectories previously obtained were grouped, and a Perron-Cluster Cluster Analysis (PCCA) was performed (Deuflhard and Weber 2005). The subspace of clustering included the whole protein volume, and the selection criteria were as previously described (Zheng et al. 2003). This resulted in 74 and 82 binding modes for charged and neutral ibuprofen, respectively. This number was further reduced by a filtering process consisting of several different metrics and manual inspection. We retained up to four binding modes for each site, and these were selected through a preliminary free energy estimation based on the Zwanzig relationship (Zwanzig 1954) (as previously described (Rocklin et al. 2013a), this can be helpful in highlighting candidate binding modes that are likely to be significantly populated when a more accurate free energy calculation is done) and the docking scores assigned by AutoDock Vina (Trott and Olson 2010). This process resulted in us retaining 18 and 13 binding modes for charged and neutral ibuprofen, respectively.

### 2.4 Absolute binding free energy calculations

A thermodynamic cycle (Boresch et al. 2003; Mobley et al. 2006, 2007) including four steps was constructed to estimate the binding affinity of ibuprofen, Δ*G*, in each of the binding modes selected. A total of six restraints (one for the distance, two for bond angles and three for dihedral torsions) were used to confine ibuprofen in the binding location (Boresch et al. 2003), and the free energy difference Δ*G*_*res*_ between a restrained and unrestrained MD run was calculated. Subsequently, ibuprofen was decoupled from the system by conducting a series of separate simulations at varying values of coupling parameter λ, ranging from 0 (full interactions) to 1 (no interactions). Coulombic interactions were decoupled first, followed by non-electrostatic ones, with softcore potentials used for Lennard-Jones interactions only. This resulted in 24 simulations carried out for 1 ns each in the presence of restraints, after which the overall Δ*G*_*site*_ term was evaluated by using MBAR (Klimovich et al. 2015; Shirts and Chodera 2008). The energetic contribution due to the presence of the restraints, Δ*G*_*unres*_, was calculated analytically (Boresch et al. 2003). The free energy of hydration of ibuprofen, Δ*G*_*water*_, was obtained by turning on again both electrostatic and non-electrostatic interactions in the presence of only solvent, in a simulation carried out for 5 ns at each λ value.

Both protonation states of ibuprofen complexed with albumin were of interest in the calculation of binding free energies. Ibuprofen ought to be deprotonated in solution because of its relatively low p*K*a (see Fig. 1), therefore for the charged state we considered the same protonation state of the ligand in solution and in the binding site. For the case of binding of neutral ibuprofen, we computed the binding free energy of taking the protonated form from solution to the binding site via the normal thermodynamic cycle, and included the additional free energy cost of taking ibuprofen from its charged form to the neutral form in solution. This additional term, which we called Δ*G*_*prot*_, is zero for charged ibuprofen and equal to –Δ*G*_*deprot*_ = 2.303 RT ln (pH – p*K*a) = 3.6 kcal/mol for neutral ibuprofen.

Finally, the binding free energy of ibuprofen for each binding mode was obtained by summing all the above contributions:

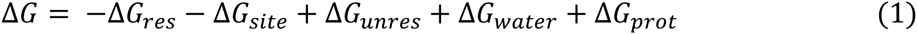

Random errors are independent for each single term of this summation, therefore the uncertainty on Δ*G* for each binding mode was calculated as the root square of the square-sum of the uncertainty of the individual contributions. In particular, uncertainties for the Δ*G*_*site*_ term were the standard error in the calculated binding free energy, as evaluated in the alchemical analysis performed through the use of MBAR after subsampling at intervals of the autocorrelation time (Shirts and Chodera 2008; Klimovich et al. 2015).

The accuracy of the calculated free energy values depends on both adequate sampling and reliability of the force field (Klimovich et al. 2015). For a host-ligand complex involving a relatively large protein such as albumin, a careful sampling of the relevant conformational states is a computational challenge, and the ability of the force field to model interactions in buried binding sites (where the ligand is well screened from the solvent) is a further cause for concern. In fact, the absence of an explicit polarizability term in classical ‘effective’ force fields reduces the dielectric response within the protein matrix and increases the strength of electrostatic interactions (Banba and Brooks 2000; Rocklin et al. 2013a). Although this effect is difficult to estimate, and might require a reparameterization of atomic partial charges to be adequately addressed, in the following we will provide some suggestions on where it could play a role in our system. The calculated binding free energies could be further influenced by finite-size effects due to the relatively small extent of the simulation box, in combination with the use of periodic boundary conditions and lattice-sum treatment of the electrostatic interactions (Rocklin et al. 2013b; Reif and Oostenbrink 2014; Lin et al. 2014). We note that this could affect the values obtained for charged ibuprofen, but has no influence on the neutral form and it is essentially independent of the binding modes examined.

## 3. Results

### 3.1 Docking predictions include a variety of possible binding sites

Initial exploration for potential binding locations of ibuprofen complexed to human serum albumin was carried out by using molecular docking. A systematic search was performed that included the whole volume for the protein, and both protonation forms for the ligand. Due to its relatively small size, ibuprofen could bind in several different internal pockets of albumin. A total of 240 binding modes were obtained, scattered in 8 different binding sites, as reported in Table 1. Two of these positions correspond to the drug sites DS1 and DS2. Other four positions are the fatty acid binding sites FA1, FA2, FA5 and FA6. The two remaining sites are in the upper and lower region of the protein cleft (hereafter abbreviated as site PC_up_ and PC_down_, respectively). Site PC_up_ partly overlaps with FA8, which is a binding location that can be occupied by capric acid but cannot fit fatty acids with a longer chain (Bhattacharya et al. 2000; Curry et al. 1998), whereas PCdown does not corresponds to any known crystallographic location for ligands bound to albumin.

**Table 1:**
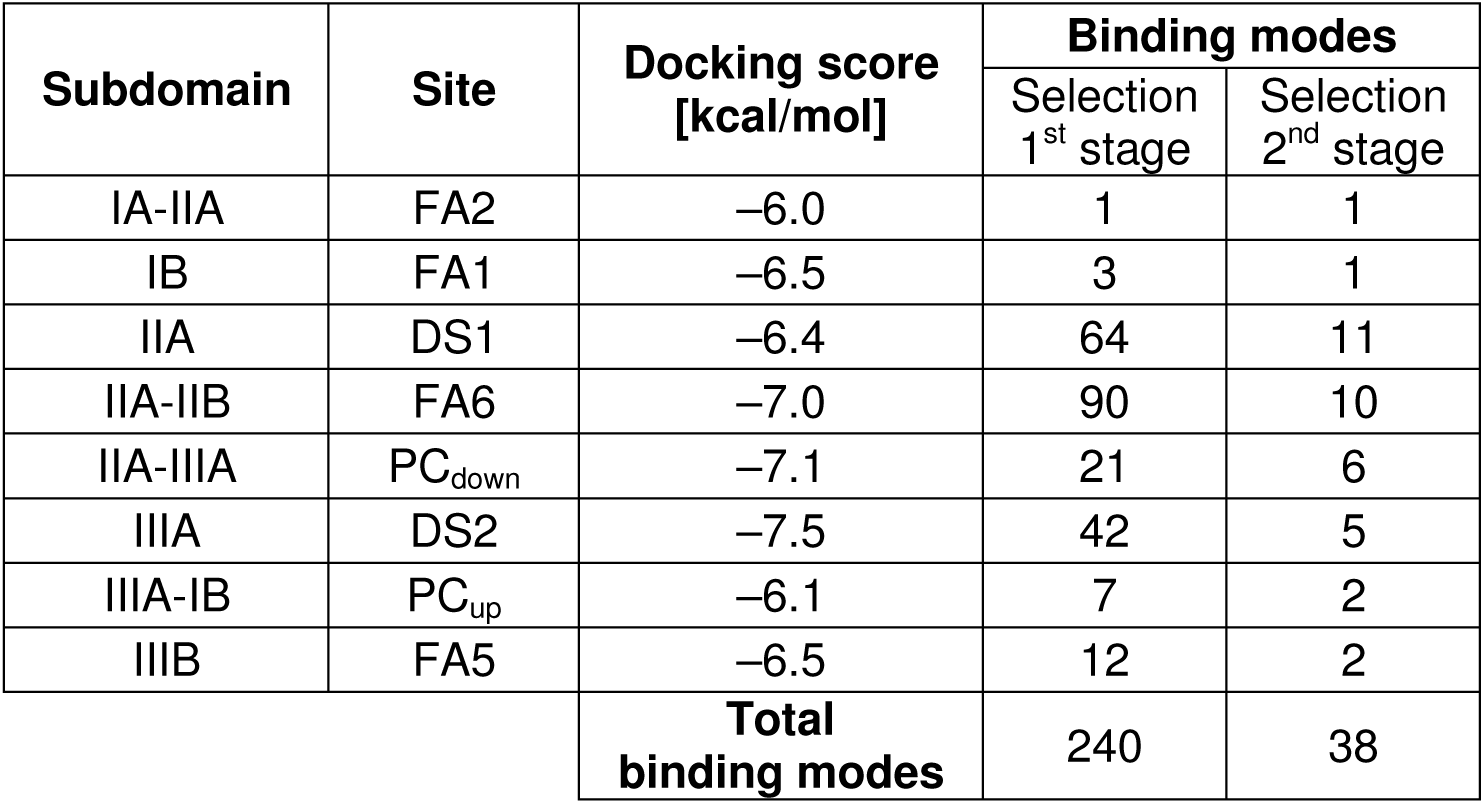
Most favorable docking score and number of binding modes found in molecular docking of ibuprofen to albumin. Selection from first to second stage was performed by a clustering procedure with a cutoff of 2.5 Å in the RMSD of atomic positions of ibuprofen.

The two crystallographic binding sites DS2 and FA6 showed two of the most favorable docking score, –7.5 and –7.0 kcal/mol, respectively. This finding is not surprising and should not be overemphasized, because in both cases ibuprofen re-docked in the binding cavity left empty upon separation of the ligand from the crystallographic complex (Ghuman et al. 2005). The other poses showed docking scores ranging from –7.0 to –6.0 kcal/mol, the two extreme values being found for ibuprofen bound in the lower and upper region of the protein cleft, respectively. Overall, the docking scores predicted on the basis of molecular docking calculations spanned an interval of only 1.5 kcal/mol, which is a difference too small to be reconciled with the variability in the ibuprofen affinity found in the experiment (Whitlam et al. 1979); furthermore, differences in the interaction energies between charged and neutral ibuprofen were exceedingly low, about 0.3 kcal/mol. These variations were of the same order of the typical uncertainties in scores provided in general by the docking technique and of the differences observed in repeating our simulation runs (about 0.1–0.3 kcal/mol in both cases).

The number of docking poses were reduced by identifying clusters of analogous binding modes differing from each other by less than 2.5 Å in the RMSD of their position. This procedure resulted in a total of 38 binding modes being selected. The number of binding modes found in each site was highly variable, ranging from 1 to 11. This quantity does not appear to be directly correlated with the volume of each protein site nor, seemingly, with other features of these cavities. The highest variety of binding modes was found for sites DS1 and FA6, suggesting that these protein pockets may accommodate ibuprofen in a multitude of different conformations.

### 3.2 MD simulations identify highly populated ligand conformations

Molecular docking is useful to identify potential binding locations, but it does not take into account the protein dynamics, which can crucially contribute to accommodate the ligand in the anchoring sites. To consider this effect, MD runs of 5 ns were carried out for the albumin-ibuprofen complex, with the drug placed in each of the 38 candidate binding modes identified by docking. Every simulation was performed with both charged or neutral ibuprofen, initially placed in the same starting position in either form, to investigate directly the consequences due to the sole difference in the protonation state. At the end of the simulations, a PCCA cluster analysis was performed (Deuflhard and Weber 2005) to select the most populated ligand conformations. This procedure resulted in 74 and 82 binding modes being identified for charged and neutral ibuprofen, respectively. Details of the distribution of the binding modes in the various protein sites are reported in Table 2.

**Table 2:**
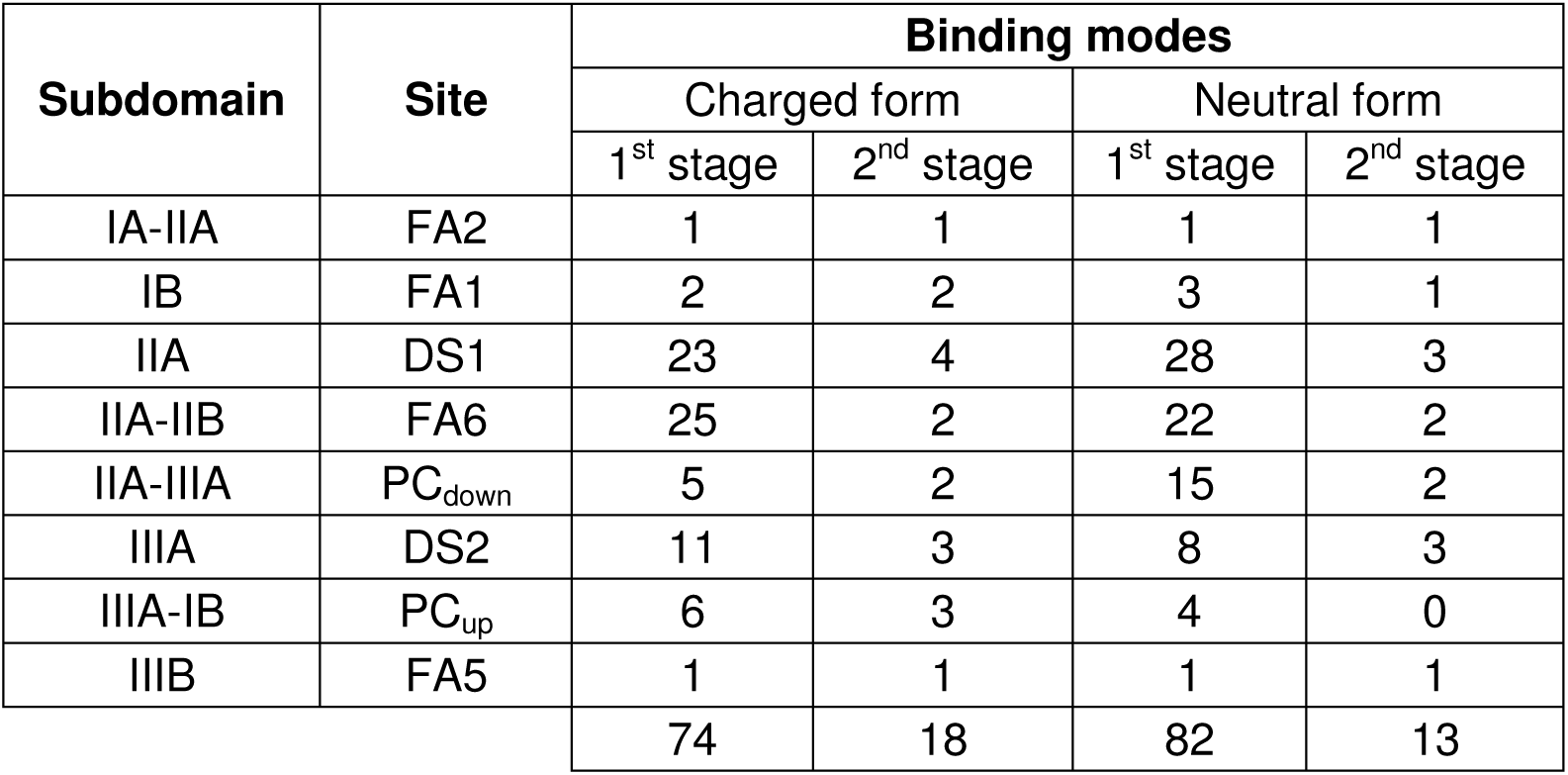
Number of binding modes found in MD simulation for ibuprofen, either in charged or neutral form, complexed with albumin. Selection from first to second stage was performed on the basis of coarse binding free energy estimates from the Zwanzig relationship (Zwanzig 1954), the number of the binding modes, and the observed population of each binding mode in our simulations.

The number of conformations was further reduced on the basis of their binding affinity, and considering a maximum number of four binding modes for each protein site. This final selection provided a total of 31 binding modes of ibuprofen, including 18 and 13 in charged and neutral form, respectively. The exact locations and conformation of the binding modes selected after the MD simulations were generally different compared to those obtained by molecular docking, although they were distributed in the same protein regions. Compared to the results previously obtained, the only possibility immediately ruled out was for ibuprofen in neutral form to associate in the upper cleft (i.e., site PC_up_) of albumin, which molecular docking had already predicted to be the least favorable region (see Table 1).

The distribution of all the possible binding modes within albumin is summarized in Fig. 2. While our process did not design for this, the resulting candidate binding modes cover all the possible ligand positions found in crystallography for any human serum albumin-ligand complex (Cuya Guizado 2014), which currently encompass 38 distinct structures deposited in the Protein Data Bank. These positions also include the possibility that ibuprofen in charged form binds in site PC_up_, which is experimentally confirmed as a binding location only for capric acid, thyroxine and iodipamide (Bhattacharya et al. 2000; Ghuman et al. 2005). In addition, a possible binding site of neutral and charged ibuprofen was considered in the lower region of the protein cleft (i.e., site PC_down_); this does not corresponds to a crystallographic location for any ligand bound to albumin, and lies roughly at the edge of site DS1. At this stage, though, all of these candidate binding modes are still considered very preliminary as they result from docking followed by relaxation via molecular dynamics.

**Fig. 2.**
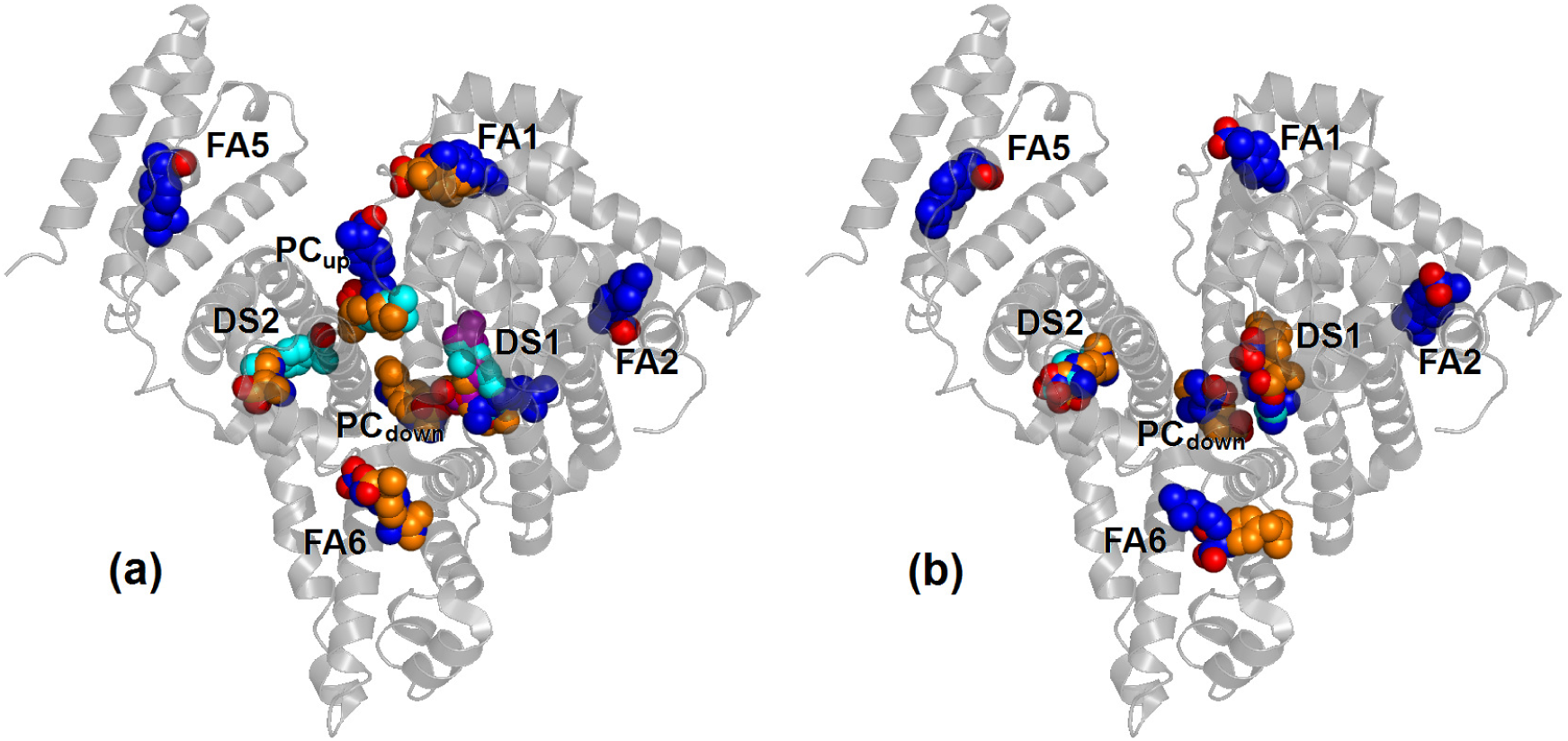
Selected binding modes of ibuprofen to human serum albumin: (a) charged and (b) neutral ibuprofen. Oxygen atoms are in red for all poses, carbons are either blue, orange, cyan or purple.

### 3.3 Binding free energies of ibuprofen span a large range of values

The predicted binding free energy of ibuprofen in each of the binding modes, determined by using alchemical free energy computations, are summarized in Table 3. A cut-off of –1.4 kcal/mol (corresponding to an association constant K_a_ < 10 M^−1^) was considered the minimum favorable energy necessary for considering a binding location to be at least a weak affinity site for ibuprofen. In 13 cases the predicted binding free energy showed values below this threshold: 4 cases for charged ibuprofen and 9 for the ligand in neutral form. These occurrences also include all the binding modes for charged ibuprofen in site PC_up_, allowing us to entirely exclude the upper region of the protein cleft as a possible binding location for ibuprofen.

**Table 3:**
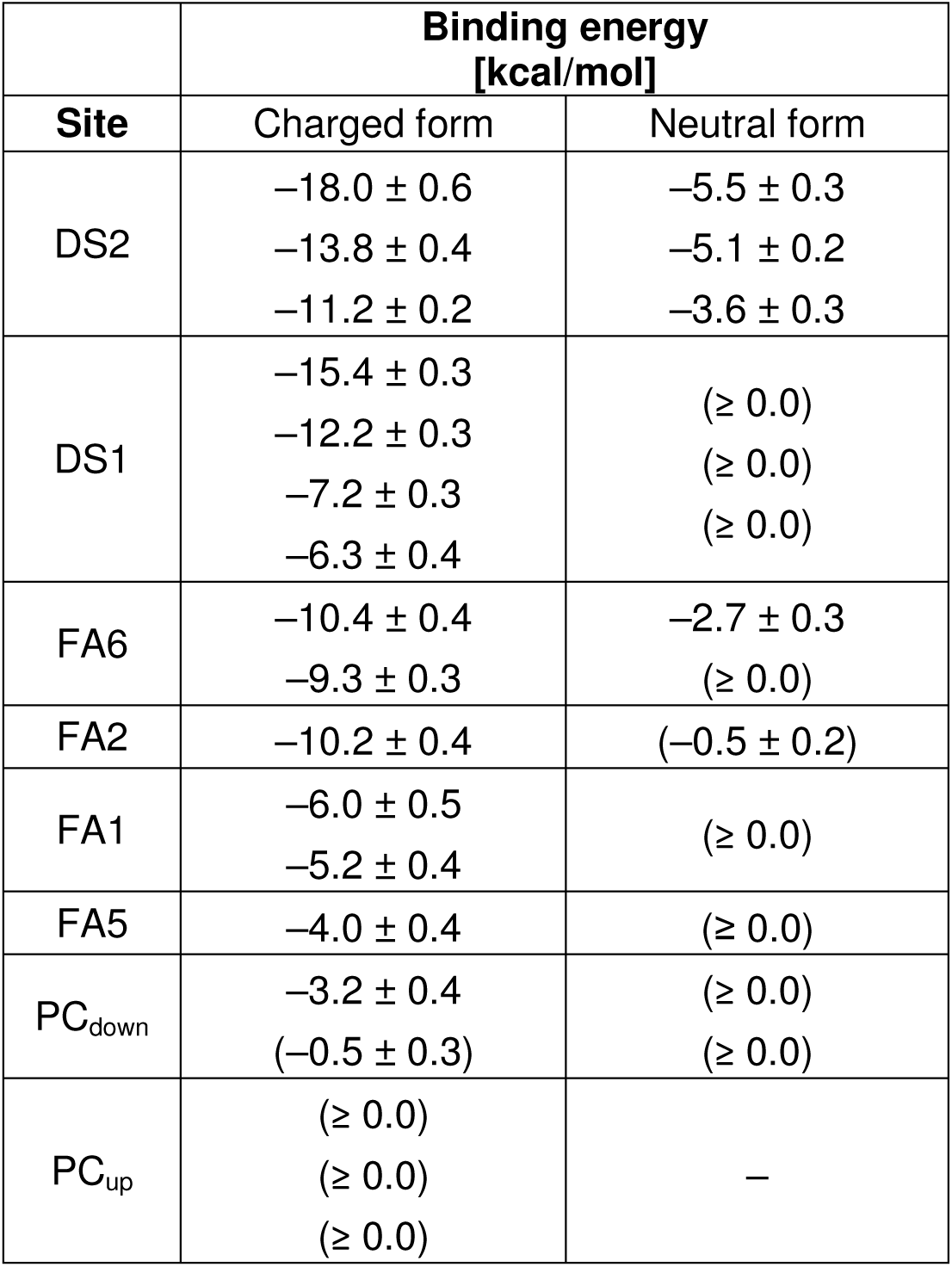
Binding energy of ibuprofen complexed with albumin determined by using alchemical free energy calculations. Free energy values ΔG_b_ > –1.4 kcal/mol are reported in parentheses, and values extending up to the positive range are indicated as ≥ 0.0 kcal/mol.

The free energy values observed are compatible with the existence of a variety of possible binding locations, which will be described (in the next two paragraphs, 3.4 and 3.5) starting with the comparison of the simulated binding modes with crystallographic data available for the albumin-ibuprofen complex (paragraph 3.4), and later analyzing the predictions obtained for the other binding sites (paragraph 3.5). Furthermore, it is clear that ibuprofen in charged form shows a much greater affinity for albumin compared to its neutral form. Overstabilization of charge-charge interactions due to inadequacies of the force field may contribute to some extent both to increase the range in the calculated values and yield overly favorable binding free energies. In addition, box-size effects could also influence the binding free energy values obtained for ibuprofen in charged form (Rocklin et al. 2013b), without modifying the relative ranking obtained for the different binding sites.

On the other hand, from a structural molecular point of view, this finding suggests that interactions between the carboxylate group of ibuprofen and positively charged (or polar) protein residues contribute significantly to determine the binding affinity. Thus, we will first analyze results observed for charged ibuprofen, and we will consider only later for comparison (in paragraph 3.6) those obtained for the neutral form. Finally, when considered separately, the ranking order in terms of affinity determined for the binding locations of charged and neutral ibuprofen are similar: site DS2 shows the highest affinity values, and less favorable binding free energies are found for some other secondary sites. This observation constitutes an additional evidence of the reliability of our alchemical free energy methodology in reproducing experimental binding locations.

### 3.4 Simulations of charged ibuprofen reproduce the crystallographic poses

The site DS2 is the most favorable binding location for ibuprofen both in the experiment and in simulation (see Table 3). For ibuprofen in its charged form, two binding modes were found with similar orientation and with binding free energies –18.0 ± 0.6 and –13.8 ± 0.4 kcal/mol, and a third one with opposite orientation and a smaller binding free energy, –11.2 ± 0.2 kcal/mol. As shown in Fig. 3, in the two binding modes of ibuprofen showing the best affinities the carboxylate group is an acceptor of hydrogen bond (HB) for both O^γ^-Ser489 (donor-acceptor distance 0.26 ± 0.01 nm) and N^ξ^-Lys414 (0.31 ± 0.01 nm). The close proximity in their locations and large difference (4.2 kcal/mol) in binding free energies both suggest that the most favorable binding mode should be drastically more populated. However, the underlying clustering procedure of the MD trajectories indicated that both binding modes (Fig. 3a and 3b) were significantly populated conformations representative of actual anchoring geometries, revealing that the range of calculated binding affinities is possibly overestimated.

Both these binding modes are very close to the location of ibuprofen detected in crystallography (represented in black in Fig. 3). One difference is that in the X-ray structure of the complex the strongest HB is formed with the hydroxyl group of Tyr411; another minor difference is that the carboxylate group of ibuprofen forms a HB with the side chain of Arg410, whereas interaction with this residue in simulation is electrostatically very favorable but does not involve the formation of a stable HB. Ibuprofen can bind within site DS2 also in an opposite orientation (Fig. 3c) compared to the crystallographic one. In this case, it can form three HBs: one with N-Gly431 (donor-acceptor distance 0.26 ± 0.01 nm) and the other two with N-Lys432 (with a distance from the two carboxylate oxygens of 0.32 ± 0.01 and 0.34 ± 0.01 nm).

**Fig. 3.**
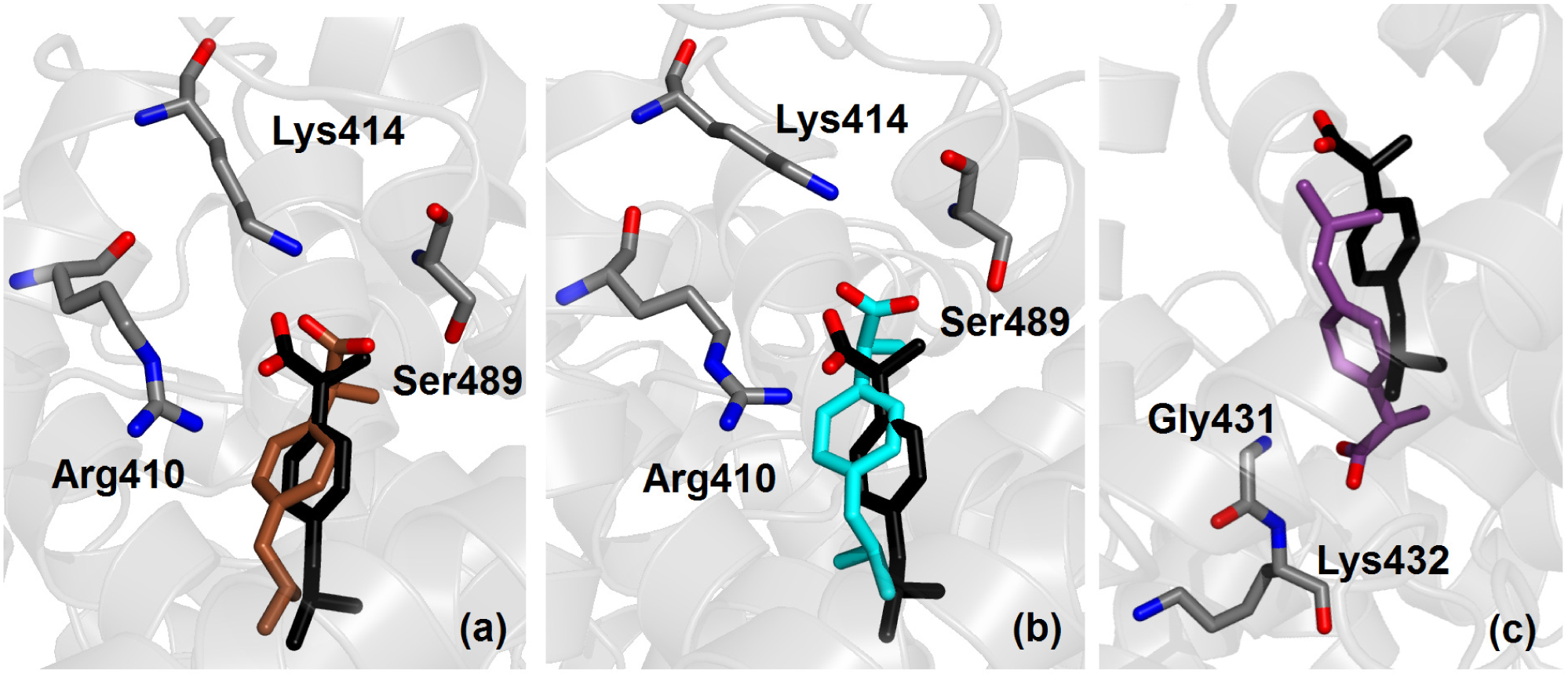
Crystallographic (black) and simulated conformation (either brown, cyan or purple) of ibuprofen in charged form bound within site DS2 of albumin. Binding modes are shown according to decreasing affinity (see Table 3) from (a) to (c). Selected protein residues are also shown; only non-hydrogen atoms are displayed.

The site FA6 is the only other binding location solved in crystallography for ibuprofen in albumin. The binding free energy calculated in simulation for the two binding modes is –10.4 ± 0.4 and –9.3 ± 0.3 kcal/mol (see Table 3), thus affinity is lower than for site DS2, but still in the nanomolar range. As shown in Fig. 4, both binding modes are very similar to the one detected in crystallography. The carboxylate group of ibuprofen interacts closely with the charged side chain of Lys351 and forms HBs with the two amine nitrogen atoms of Leu481 and Val482. For the favorite binding mode, donor-acceptor distance is, respectively, 0.29 ± 0.01 and 0.30 ± 0.01 nm for the two HB.

**Fig. 4.**
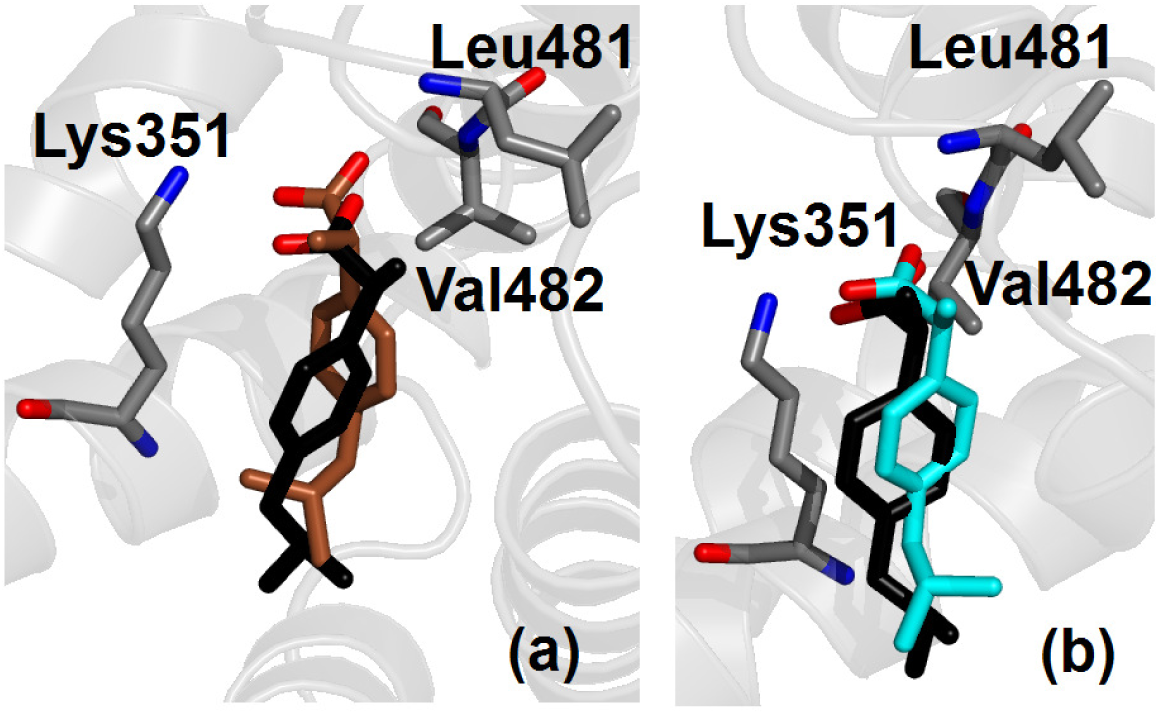
Crystallographic (black) and simulated conformation (either brown or cyan) of ibuprofen in charged form bound within site FA6 of albumin. Binding modes are shown according to decreasing affinity (see Table 3) from (a) to (b). Selected protein residues are also shown; only non-hydrogen atoms are displayed.

### 3.5 Several additional binding sites are predicted for ibuprofen

The simulation results show that site DS1 is the second most favorable binding site for charged ibuprofen interacting with albumin. The binding conformations obtained in simulation are shown in Fig. 5. Two binding modes were detected with highly favorable energies, –15.4 ± 0.3 and –12.2 ± 0.3 kcal/mol, and other two with lower affinity, –7.2 ± 0.3 and –6.3 ± 0.4 kcal/mol (see Table 3). Ibuprofen forms HBs with N^ξ^-Lys199, N^η^-Arg218 and N^η^-Arg222, with an average donor-acceptor distance of 0.28 ± 0.01 nm in the first two cases and 0.31 ± 0.01 nm in the third one; exceptions are the second most favorable binding mode (Fig. 5b), which does not form a HB with N^η^-Arg222 but it does with N^ξ^-Lys195 (with distance 0.31 ± 0.01 nm), and the third most favorable binding mode (Fig. 5c), which does not form a HB with Arg218. Therefore, although the two binding modes with the most favorable free energies should be significantly preferred, overall ibuprofen in charged form shows a variety of possibilities to interact with albumin in site DS1. A direct comparison with crystallography is not possible because an accurate experimental position of the binding pose was not reported, although indications were found of the presence of the ligand in this site (Ghuman et al. 2005).

**Fig. 5.**
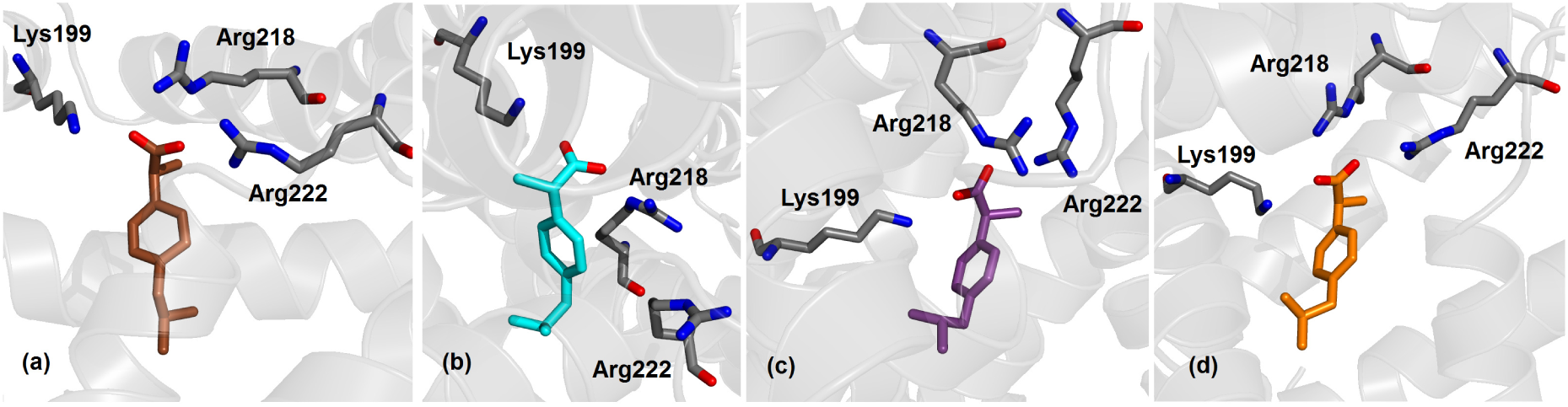
Ibuprofen in charged form bound within site DS1 of albumin; binding modes are shown according to decreasing affinity (see Table 3) from (a) to (d). Selected protein residues are also shown; only non-hydrogen atoms are displayed.

The binding affinity of site FA2 is –10.2 ± 0.4 kcal/mol (see Table 3) and is comparable with the one of site FA6, in agreement with the indications of experiments (Fanali et al. 2012), although only the latter has been detected in crystallography. As shown in Fig. 6, ibuprofen in site FA2 forms a HB with the side chain of Arg10, with a donor-acceptor distance of 0.32 ± 0.02 nm. The other three possible binding sites investigated (FA1, FA5 and PC_down_) show low affinity values, indicating that they have lower specificity for ibuprofen and could be occupied only when all the other locations are already saturated. In particular, binding free energy site for site PC_down_ is very low, indicating that this site has low specificity for ibuprofen.

**Fig. 6.**
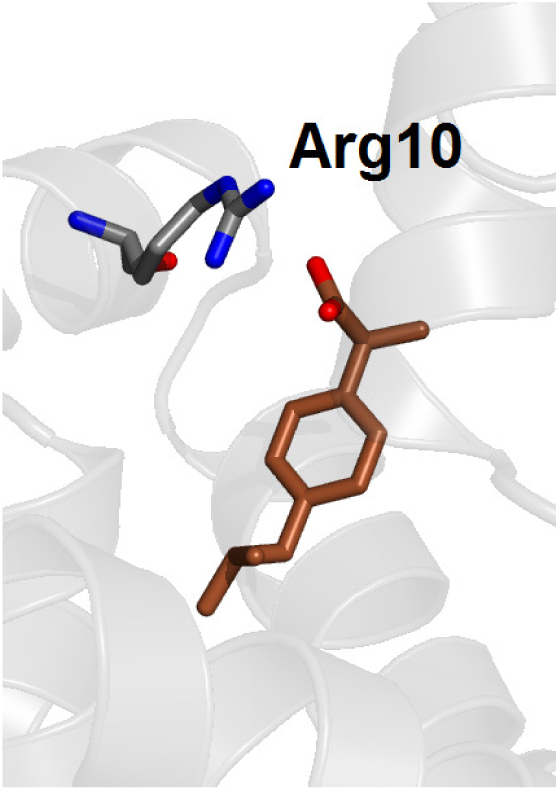
Ibuprofen in charged form bound within site FA2 of albumin. The hydrogen-bonding residue Arg10 is also shown; only non-hydrogen atoms are displayed.

### 3.6 Ibuprofen binds albumin only in charged form, not neutral

The binding of ibuprofen in neutral form to albumin may appear to have a significant affinity in absolute terms, because the values obtained in simulation extend up to –5.5 kcal/mol for site DS2 (see Table 3), i.e. to micromolar affinity. Nevertheless, binding free energy values in every binding location are systematically more favorable for ibuprofen in charged form compared to the neutral one, even when the best binding geometry found for the two protonation states is almost identical. This evidence seems to exclude the possibility that ibuprofen binds significantly to albumin in neutral form.

An example is shown in Fig. 7, where the binding mode with the most favorable free energy for neutral ibuprofen in site DS2 is reported. This binding mode is close to the one observed in crystallography (in black in Fig. 7), and similar to the one already found for charged ibuprofen (see Fig. 3), which has however a binding free energy predicted to be more favorable by 12.5 kcal/mol (see Table 3). The difference in the affinity between the two protonation states of ibuprofen is essentially due to electrostatic interactions that are missing in the neutral state. Other contributions, such as the possibility for neutral ibuprofen to form a HB between its unprotonated O atom and N^ξ^Lys414 (see Fig. 7), only play a minor role. The same is observed also for binding modes in other protein sites, which have less favorable interactions with the protein due to the absence of significant electrostatic contributions, and show only low affinities values. In particular, for site PC_down_ the calculated value of binding affinity is particularly unfavorable, ruling out this location as a binding site for ibuprofen in any form of protonation.

**Fig. 7.**
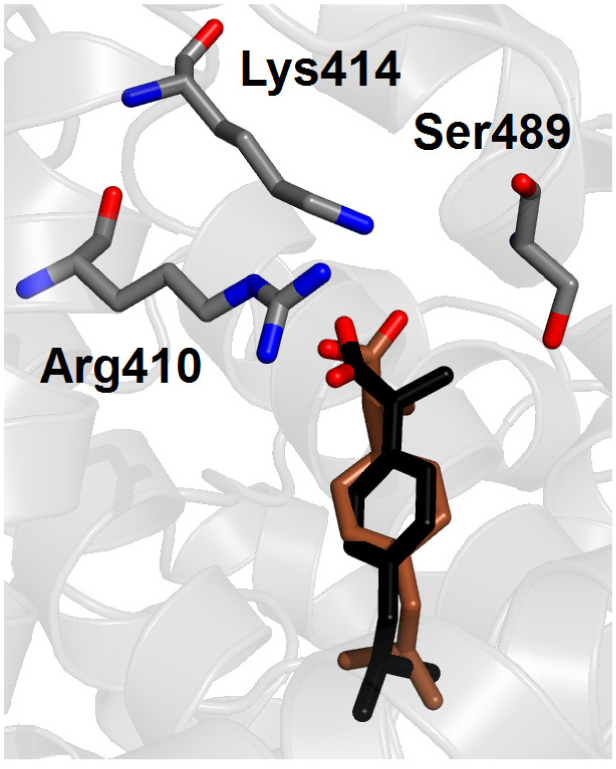
Crystallographic (black) and simulated conformations of the most favorable binding mode (brown) of ibuprofen in neutral form in site DS2 of albumin. Selected protein side chains are also shown; only non-hydrogen atoms are displayed

It is interesting to note that the minimum difference in binding free energy between the two protonation states of ibuprofen in the same binding location is found for sites FA6, which is a shallow trench on the protein surface, thus relatively open and accessible to the solvent. In contrast, binding free energy differences between charged and neutral form of ibuprofen are very large for both Sudlow’s drug sites, which are secluded inner cavities and are predominantly apolar, except for the additional presence of either two patches of basic and polar residues (site DS1) or a single polar patch (DS2). This observation supports the idea that there may be an inadequate environmental polarization in the most solvent-sheltered protein regions that would have an impact on the calculated interaction energies, systematically affecting the values obtained for charged ibuprofen over those relative to the neutral form.

## 4. Discussion

Mapping the exact location and binding affinity of ligands to human serum albumin is remarkably important in pharmacology, because it contributes to determine the adsorption of currently administered drugs and could help in the design new active compounds. This task is difficult, both from a theoretical and experimental point of view, for a number of reasons. The most important one is that albumin possesses several inner pockets, variously accessible to different ligands (Ghuman et al. 2005): e.g., seven sites for long-chain fatty acids, five for thyroxine, etc. In addition, multiple binding modes can sometimes be found in a single binding site; e.g., stearic acid can bind in two configurations with opposite direction in site FA2 (Bhattacharya et al. 2000). Some compounds are also titratable, and their binding free energy is affected by their protonation state.

Ibuprofen is an important molecule from a pharmaceutical point of view, and it can be considered a model ligand for a systematic *in silico* investigation of the binding properties of albumin. In fact, several experimental techniques suggest that ibuprofen has multiple binding locations (Kosa et al. 1997), and some of them can possibly be occupied in more than a single binding mode. Furthermore, the protonation state of this ligand may be either charged or neutral, although the former state is more likely to occur at physiological pH. Both protonation forms were simulated and the results were compared.

Our investigation on the possible binding locations and affinity for ibuprofen complexed to albumin has relied on a combination of simulation methods. Molecular docking was used as an ancillary technique for a first screening of the protein surface, in search for binding cavities to be (potentially) occupied by ibuprofen. The positions determined included all of the binding pockets already known in crystallography for ligands bound to albumin (Cuya Guizado 2014), plus a supplementary location in the lower region of the central protein cleft. The binding modes obtained were subsequently refined through MD simulations that allowed the protein to accommodate the ligand. This procedure let ibuprofen adapt its position and sample the most populated and energetically favorable conformations, and alchemical free energy calculations were finally used to evaluate the binding free energies in these positions.

Several arguments strongly suggest that our predicted binding modes may provide a realistic molecular picture. First, the most favorable binding modes reproduce very accurately the position found in crystallography for ibuprofen bound to human serum albumin (Ghuman et al. 2005). Second, the ranking of the binding free energies calculated in the different positions is in excellent agreement with the indications of experiments (Fanali et al. 2012). Third, simulation of ibuprofen in neutral form does not alter significantly such ranking, and yet the binding affinity drastically reduces compared to the charged state, ruling out the possibility that the ligand is protonated within the protein matrix – as it was reasonable to expect. Finally, although no additional binding site compared to the ones known for other ligands was found for ibuprofen within albumin, a multiplicity of possible binding locations and modes were found in simulation, suggesting an immediate rationale to explain some previously discordant experimental observations on the exact number of such locations and their respective affinity (Itoh et al. 1997; Kosa et al. 1997; Montero et al. 1990).

A direct comparison between the affinity values found in simulation and the experimental ones is not straightforward. For instance, experiments are often performed under restricted conditions, e.g. with albumin solved in a buffer or immobilized on a substrate. For this reason, estimates of the binding constants of ibuprofen to albumin may be at variance even in different ‘wet lab’ experiments (Kosa et al. 1997), and it is not surprising that simulation results obtained under ideal, unperturbed conditions may further differ. Conversely, computational methods are also not immune from systematic errors that may affect the correct estimate of the binding free energy. As an example, discrepancies up to 4.5 kcal/mol with the experiment were reported in the affinity of fatty acids calculated by using molecular mechanics Poisson–Boltzmann surface area (MM-PBSA), on of the most accurate method applied to estimate ligand binding to human serum albumin to date (Fujiwara and Amisaki 2008).

In our case, typical unavoidable difficulties associated with MD simulations, such as intrinsic limits in both sampling and force field, might affect the quantitative estimates of the determined free energy values. We note that finite-size effects (Rocklin et al. 2013b) may influence the results obtained for charged ibuprofen, without affecting those for the neutral form. However, these effects are essentially independent of the specific binding location, thus would not alter the overall ranking of the preferred binding sites. Therefore, we point out to an excess in the stabilization of charge-charge interactions as the possible predominant source of error in our calculations, in close similarity with work on cytochrome *c* peroxidase (Banba and Brooks 2000; Rocklin et al. 2013a). Although not easy to prove in the practice, this hypothesis would explain in a rather straightforward way three systematic effects that are evident in our data: (i) overly favorable binding free energies, particularly for (but not limited to) charged ibuprofen; (ii) large variations in the calculated values across all the possible binding sites for the charged ligand form; and, (iii) differences between binding energies of protonated versus deprotonated ibuprofen in the same binding site are much greater for the most internal and apolar binding pockets (i.e., site DS1 and DS2), where the solvent cannot provide adequate screening of electrostatic interactions.

These effects are particularly evident for site DS2, which is the most favorable binding location found for ibuprofen – in agreement with experiment. For comparison, binding free energies varying in the range between –8.6.and –6.9 kcal/mol for S–ibuprofen in the highest affinity site of albumin were previously reported in experiments performed with several techniques, which included ultrafiltration after *in vivo* administration of ^3^H-labeled ibuprofen (Paliwal et al. 1993), microcalorimetry (Cheruvallath et al. 1996), circular dichroism (Cheruvallath et al. 1997), and high-performance liquid chromatography (HPLC) either by equilibrium dialysis (Rahman et al. 1993), or on an immobilized albumin column (Hage et al. 1995), or using a chiral fluorescent derivatizing reagent (Itoh et al. 1997).

The second most favorable binding site found in simulation is DS1, and shows several possible binding modes that may contribute to explain a number of puzzling experimental observations. In fact, ibuprofen was reported (Kosa et al. 1997) to be indifferent to the presence of warfarin and prone to an anti-cooperative behavior towards phenylbutazone, both typical marker ligands of site DS1. In contrast, warfarin is displaced from this site in haem-bound albumin under excess of ibuprofen, after the latter has saturated its primary site DS2 (Baroni et al. 2001); the same effect was observed for unliganded albumin after sequential addition of ochratoxin A and ibuprofen (Il'ichev et al. 2002). A strong interference was also found in the concurrent association of quercetin, which is likely to bind primarily in site DS1 (Dufour and Dangles 2005). These competitive behaviors were explained by assuming that ibuprofen binds preferentially to albumin in site DS2, with DS1 available as a secondary site. Nevertheless, Hage and coworkers (Hage et al. 1995) reported the presence of two high affinity sites with similar binding free energy for S– ibuprofen bound to albumin. Their experiment were performed on immobilized albumin, and this rationale is usually given to account for differences with other studies (Itoh et al. 1997). However, a number of high affinity sites in between 1 and 2 was observed to be necessary to fit experimental observations for both ibuprofen isomers also by Rahman and coworkers (Rahman et al. 1993). Thus, DS1 may be more appropriately described as a versatile high-affinity binding site for ibuprofen, as our simulation results readily suggest.

The multiplicity of possible binding modes in site DS1 would also explain why the exact position of ibuprofen in this site is poorly solved in the crystallographic complex (Ghuman et al. 2005) compared with site FA6, although the latter is found only as a secondary site both in other spectroscopic experiments (di Masi et al. 2011) and in our free energy calculations. The higher adaptability of site DS1 in lodging ibuprofen compared to DS2 might also justify why only the former shows high affinity for the (biologically less relevant) R– isomer (Hage et al. 1995), although this hypothesis should be further tested.

Other binding locations, with comparatively less favorable affinity, were found in simulation in sites FA6 and FA2. Site FA6 has two binding modes, with most the favorable one showing a binding geometry in close agreement with crystallographic data (Ghuman et al. 2005). In contrast, site FA2 has a single binding mode and was not found in X-ray experiments, but it was already reported as an experimental binding location for ibuprofen (di Masi et al. 2011); in this case, the results of our MD simulations allow us to predict details of the binding geometry that were formerly unknown. Finally, sites FA1 and FA5 are low affinity binding locations that could constitute secondary association sites at sufficiently high concentrations of ibuprofen, and even the central cleft of albumin (site PC_down_) may bind ibuprofen under opportune conditions, although with very low affinity.

The existence of multiple binding sites for ibuprofen was already hypothesized (Dufour and Dangles 2005) to explain the increase of fluorescence quenching in flavonoids/albumin complexes taking place upon addition of relatively high concentrations (ibuprofen/albumin molar ratio > 4). Up to 6–10 binding modes where also necessary in former studies to explain some experimental data (Kosa et al. 1997; Montero et al. 1990; Whitlam et al. 1979). Such an assortment of possibilities may be relevant *in vivo* because, although at typical therapeutic concentrations (~ 5⋅10^−5^ M, or 10 mg/L) the molar ratio ibuprofen/albumin is about 0.1 (Itoh et al. 1997), other endogenous and exogenous ligands compete at the same time to occupy the binding sites of the protein. In particular, bulkier drugs generally have no other possibilities than binding in Sudlow’s DS1 and DS2 sites. Thus, the variety of binding possibilities found in our simulations reveal the versatility and promiscuity of albumin in forming a complex with a multiplicity of ligands, which is one of the major keys in determining the importance of this transport protein in the blood.

## 5. Conclusion

The binding of ibuprofen to human serum albumin is important from a pharmacological point of view, and constitutes a relevant test case to assess the general ability of MD simulations to reproduce experimental findings and predict binding locations in this essential plasmatic protein. Alchemical free energy calculations were found to accurately model both the geometry and the binding affinity of ibuprofen in the various protein binding sites. The simulation results explain several findings obtained by using different spectroscopic techniques, and make testable predictions that will help to complete an accurate description of the binding sites of human serum albumin. The simulation methods here described could also be immediately extended to investigate the binding of other endogenous and exogenous molecules to albumin, or to other drug-binding proteins with an equally complex structural architecture and dynamical behavior. More ambitiously, this work may constitute a starting point for more advanced simulations, to investigate interactions among competing ligands with the same host macromolecule, or to unravel the role of allosteric regulation in the binding of small molecules to transport proteins.

## Abbreviations

DS1/DS2: Sudlow’s drug site 1/2
FA1/7: fatty acid site 1/7
HB: hydrogen bond
HPLC: high-performance liquid chromatography
MD: molecular dynamics
MM-PBSA: molecular mechanics Poisson–Boltzmann surface area
PCCA: Perron-cluster cluster analysis
PC_down_/PC_up_: lower/upper protein cleft site
PME: Particle-Mesh Ewald
RMSD: root mean square deviation.

## Acknowledgments

DLM appreciates financial support from the National Institutes of Health (1R01GM10888901).

